# CASTLE: Cell-type Aware SpaTial domain detection via contrastive Learning Embedding

**DOI:** 10.64898/2026.02.12.705478

**Authors:** Aoqi Xie, Yuehua Cui

**Affiliations:** Department of Statistics and Probability, Michigan State University, East Lansing, MI

**Keywords:** Spatial clustering, Graph convolution network, Cell type composition, Contrastive learning

## Abstract

With advances in spatially resolved transcriptomics across platforms and resolutions, it is now possible to measure gene-expression profiles while preserving the tissue microenvironment. Spatial clustering is central to these analyses, and recent graph neural network (GNN)-based approaches have greatly improved their accuracy. Nevertheless, precisely delineating spatial domain boundaries remains a significant challenge. Here we present a novel method CASTLE (Cell-type Aware SpaTial domain detection via contrastive Learning Embedding) by leveraging cell type information for improved spatial domain detection. CASTLE first integrates spatial proximity to construct a spatial graph, refined from cell-type similarity or expression similarity when cell type information is unavailable. Then, a self-supervised, local-context contrastive objective aligns embeddings with their immediate microenvironments. Evaluated across multiple tissues and technology resolutions, CASTLE outperforms state-of-the-art methods and detects more accurate and stable spatial domains. In downstream analyses, it resolves fine-grained tissue structures and differentiates functional subtypes, supporting more nuanced biological interpretation.

## 1 Introduction

Spatially resolved transcriptomics (ST) includes positional context to gene expression data and enables analyses that are impossible with dissociated scRNA-seq alone, including spatial domain delineation, discovery of spatially variable genes, inference of cell–cell communication, and tumor microenvironment profiling[1, 2, 3, 4]. Among downstream tasks, spatial domain (region) identification is foundational. Inaccurate domains can lead to poor performance for domain-specific marker gene detection, and spatial microenvironment characterization[5, 6].

A variety of strategies have been developed for spatial domain identification. Traditional clustering (Louvain[7], Leiden[8]) is widely used in Scanpy[7] and Seurat[9] pipelines as part of integrated analyses. Probabilistic models formalize spatial smoothness via Markov random fields and neighborhood-structured priors, for example, Giotto[10] implements a hidden Markov random field (HMRF) to detect spatially coherent domains, and BayesSpace[11] employs a fully Bayesian framework with neighborhood priors to enhance resolution and improve clustering. Recently developed spatial clustering methods often employ graph neural networks (GNNs), which integrate information from spatial graphs to learn embeddings that capture both gene expression and spatial context, resulting in more comprehensive and robust analyses. Among the various recently developed methods for spatial domain detection, differences arise in how they acquire information to construct low-dimensional embeddings for subsequent domain identification. For example, SpaGCN[12] integrates gene expression, spatial coordinates and histology image within a GCN model; stLearn[13] performs morphology-aware normalization and integrates spatial location with image-derived features before graph-based clustering; DeepST[14] pairs a denoising autoencoder with a GNN autoencoder to learn joint embeddings from expression, spatial location and histology image; SEDR[15] couples a deep autoencoder for expression with a variational graph autoencoder for spatial structure to yield unified latent representations; STAGATE[16] employs a graph attention autoencoder whose adaptive attention captures boundary similarity and optionally incorporates a cell type–aware module; CCST[17] implements a GCN-based unsupervised clustering framework built on deep graph infomax (DGI) objectives; SpaceFlow[18] adopts a contrastive learning strategy by generating negatives via spatial-graph permutation during encoder training; GraphST[19] applies augmentation-based self-supervised contrastive learning to encode both expression similarity and spatial proximity; and stAA[20] applies an adversarially regularized variational graph autoencoder that refines embeddings with a Wasserstein adversarial objective using pre-clustering labels.

Despite strong use of spatial neighborhood graphs, most methods do not explicitly and systematically model the influence of local cell-type composition. In many tissues, the spatial distribution of cell types is a primary determinant of domain structure[21, 22, 23]. For example, in the human dorsolateral prefrontal cortex (DLPFC), laminar domains are defined by graded mixtures of excitatory neuron subtypes, interneurons, astrocytes, oligodendrocytes, and microglia, yielding sharp layer-to-layer composition shifts that are reproducible across sections and donors[24]. In the mouse main olfactory bulb (MOB) data, distinct cell populations, including periglomerular and tufted cells in the glomerular layer, mitral cells in the mitral cell layer (MCL), and granule cells in the glomerular layer (GL), form concentric and well-separated strata.[25, 26]. Similarly, the hippocampal region displays distinct cellular mosaics formed by CA1 and CA3 pyramidal neurons and dentate gyrus granule cells[27]. At single-cell resolution, MERFISH[28] and STARmap[29] atlases map these molecularly defined cell classes into sharply delineated layers and niches, further underscoring that composition encodes spatial domains rather than merely correlating with them[3, 4]. Moreover, cell type deconvolution studies (e.g., RCTD[30], SPOTlight[31], cell2location[32]) consistently reveal structured spatial gradients and patches of cell-type mixtures, and these compositional maps align with histological boundaries and microenvironmental niches. Taken together, these observations argue that explicitly modeling cell-type composition and cell-type-aware similarity in the graph could yield more faithful neighborhoods, sharper domain boundaries, and more reliable downstream inference than proximity alone.

While some methods hint at cell-type awareness, they do not incorporate it into graph construction in a principled way. For example, STAGATE[16]’s optional “cell type–aware” spatial network is obtained by running the Louvain algorithm on low-dimensional embeddings based on gene expressions (e.g., principal components) to assign a predefined cluster label per spot for multicellular data (e.g., 10x Visium). It then uses those labels (i.e., pre-trained domain labels) to adjust the neighborhood graph to further detect spatial domains. This process introduces a form of label feedback, or circularity. That is, cluster assignments derived from the same features are subsequently used to reshape the graph for further cluster detection. Such feedback can reinforce initial classification errors and obscure genuine boundaries that span multiple preclusters. It may also artificially inflate apparent performance in regions where spot composition is inherently heterogeneous.

**CASTLE** (Cell-type Aware SpaTial domain detection via contrastive Learning Embedding) addresses these gaps by integrating spatial proximity and cell-type information directly into graph construction, using deconvolved compositions for spot-level data or cell-level annotations for single-cell platforms, with an expression-based fallback when labels are uncertain or unavailable. A self-supervised, local-context contrastive objective then learns embeddings aligned to each spot/cell’s microenvironment, while a lightweight encoder-decoder reconstructs expression to regularize the representation. Benchmarks across 10x Visium[33], classic ST[25], MERFISH[28], STARmap[29], Stereo-seq[34], and Slide-seqV2[35] in diverse tissues (human brain and breast cancer; mouse brain, hippocampus, and MOB) show that CASTLE achieves higher clustering accuracy and boundary fidelity than state-of-the-art counterparts such as GraphST, STAGATE and stAA, highlighting the value of cell-type–aware graph design for robust spatial domain identification.

## 2 Methods

### 2.1 Overview of CASTLE and data pre-processing

For all datasets, pixels outside the main tissue area were removed. Raw counts were normalized per spot by library size and then log-transformed. Gene-expression features were scaled to zero mean and unit variance. We selected the top 3,000 highly variable genes (HVGs) and used this matrix as input to CASTLE. Unless noted otherwise, the same pre-processing was applied uniformly across all methods for fair comparison. Figure 1 shows the overview of the CASTLE algorithm.

**Figure 1:**
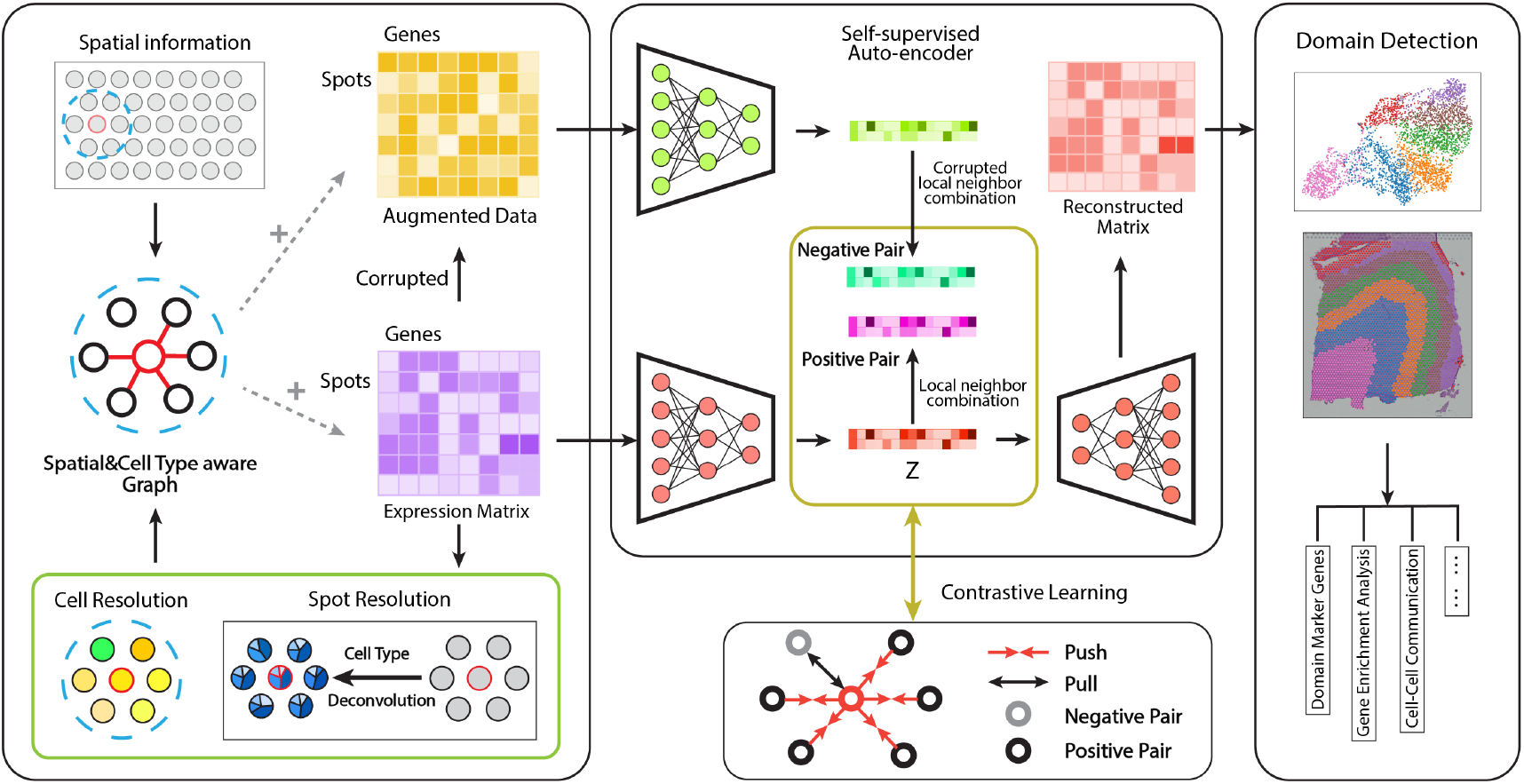
Overview of CASTLE. CASTLE starts with the gene-expression matrix and performs data augmentation for self-supervised contrastive learning. It builds a cell type–aware spatial graph by combining spatial proximity with biologically informed similarity from (deconvolved) cell-type compositions or expression-based similarity when labels are unavailable. Using this graph, CASTLE learns low-dimensional embeddings that preserve informative signals from gene expression, spatial location, and the local neighborhood context. A decoder then maps these embeddings back to the original feature space to reconstruct the expression matrix, which is then used for spatial clustering and supports downstream analyses such as domain interpretation, pathway enrichment, and microenvironment profiling.

### 2.2 Cell type-aware spatial graph construction

#### Spatial neighborhood graph

We encode spatial context as an undirected *k*-nearest neighbor graph *G* = (*V, E*) over *n* pixels. Let *A* ∈ ℝ^*n*×*n*^ denote the adjacency matrix. Pixels *i* and *j* are connected if one lies among the *k* nearest neighbors of the other in spatial location; edges are symmetrized to ensure an undirected graph.

#### Cell-type similarity

Each measurement location (pixel) is treated uniformly, with methodological adjustments made to account for platform-specific characteristics.

- **With spot-level resolution data (e.g., 10x Visium):** Estimate a cell-type composition vector per spot via cell type deconvolution; define cell-type similarity between two spots with the cosine similarity. Users can apply a reference-based deconvolution method such as RCTD[36] or reference-free methods such as gwSPADE[37] when reference information is unavailable.
- **With single-cell resolution data (e.g., MERFISH):** Conduct cell type annotations to get cell type labels for all cells; set similarity to 1 for the same cell type and 0 otherwise (or a soft similarity if subtype labels are available).
- **With no labeled or subcellular resolution data:** Compute a label-free similarity from gene expressions by taking the cosine similarity over low-dimensional embeddings.

#### Cell type-aware spatial graph with edge weighting

For a candidate edge (*i, j*), we refine spatial proximity with cell type or expression similarity. Let *s*_*ij*_ denote the similarity computed as above (composition cosine, same-type indicator, or PCA-cosine), and let *τ* be a minimum threshold. We define

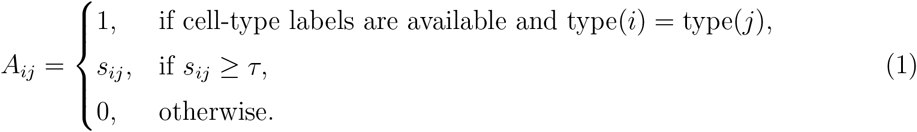

We choose *k* and *τ* so that each pixel has approximately 3-6 neighbors on average.

Incorporating cell-type information (via deconvolved compositions or single-cell labels) enriches the neighborhood graph with biological similarity, reducing spurious cross-type links that arise from spatial proximity alone. When labels are unavailable or noisy, the PCA-cosine fallback maintains robustness by capturing expression-level similarity. The resulting graph better reflects local tissue organization and guides CASTLE to learn representations consistent with underlying spatial structure.

### 2.3 Self-supervised auto-encoder

Like GraphST[19], CASTLE incorporates both data augmentation and self-supervised learning.

#### 2.3.1 Data augmentation

##### Feature-permutation augmentation (corrupted view)

Given the spatial neighbor-hood graph *G* = (*V, E*) with |*V*| = *n* pixels and the normalized gene-expression matrix *X* ∈ ℝ^*n*×*p*^ with *p* genes, we construct a corrupted graph by permuting features across pixels while preserving geometry. Let *π* be a random permutation of {1, …, *n*} and denote the permuted feature entry as:

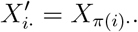

We keep the node set and geometric neighborhoods fixed (same *V* and spatial *k*-NN structure), and recompute edge weights from *X*^′^ (and update edges if similarity-thresholded) to obtain the corrupted graph

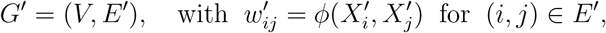

where *ϕ*(·, ·) denotes the chosen feature-similarity function (e.g., cosine on compositions or PCA embeddings on gene expressions). Thus, *G*^′^ preserves the spatial topology of *G* but disrupts local feature coherence, providing a hard negative view for contrastive learning. Based on *G*^′^, we generate a corrupted adjacency matrix *A*^′^.

#### 2.3.2 GCN-based auto-encoder

We design a single-layer graph convolution network (GCN) encoder to learn pixel representations that capture informative gene expression and spatial signals. Given the spatial graph *G* = (*V, E*) with adjacency *A* and the normalized expression matrix *X*, the encoder aggregates neighborhood information as

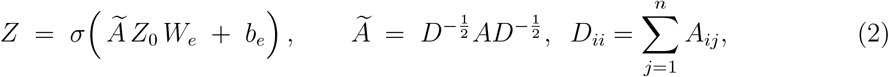

where *Z*_0_ is set as the original input gene expressions *X. W*_*e*_ and *b*_*e*_ are trainable weights and bias, respectively, and *σ* is a pointwise nonlinear activation function (e.g., ReLU). Here, *Z* denotes the latent spot representation produced by the encoder.

The decoder mirrors the encoder to reconstruct gene expression from *Z*:

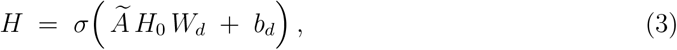

where *H*_0_ is set as the output representation *Z. W*_*d*_ and *b*_*d*_ are trainable decoder parameters and *H* ∈ ℝ^*n*×*p*^ is the reconstructed expression matrix.

To exploit full expression information, we train the model by minimizing a self-reconstruction loss,

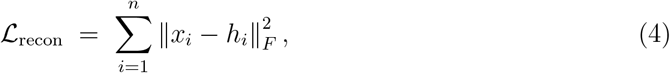

where *x*_*i*_ and *h*_*i*_ denote the *i*-th rows of *X* and *H*, respectively (i.e., the original normalized and reconstructed gene-expression profiles of pixel *i*). Bias vectors are broadcast across rows, and the same normalized adjacency *Ã* is used in both encoder and decoder.

#### 2.3.3 Self-supervised contrastive learning on local contexts

To make the reconstructed outputs *H* more informative and discriminative, we adopt a self-supervised contrastive learning (SCL) strategy that encourages embeddings to capture the *local spatial context* of each spot[19]. Given the original graph *G* = (*V, E*) with features *X* and the corrupted view *G*^′^ = (*V, E*^′^) with permuted features *X*^′^, the GNN encoder produces two representation matrices *Z* and *Z*^′^. For each pixel *i*, we summarize its immediate neighborhood as a local-context vector by averaging the embeddings of its adjacent nodes and passing them through a nonlinear sigmoid function:

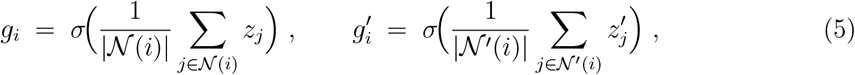

where 𝒩 (*i*) denotes the neighbors of the original pixel *i*, 𝒩^′^(*i*) denotes the neighbors of corrupted pixel *i*. Unlike global readouts used in DGI[38], this readout focuses on *immediate* neighbors, reflecting the spot’s neighborhood microenvironment.

We treat (*g*_*i*_, *z*_*i*_) as a positive pair and 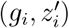 as a negative pair. Let Φ (·, ·) : ℝ^*d*^ × ℝ^*d*^ → ℝ be a discriminator that scores pairs (higher for positives). Using a binary cross-entropy objective, the contrastive loss on the original view is

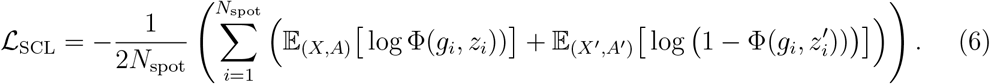

Because *G*^′^ shares the similar topology structure with *G*, we also define a symmetric loss that swaps the roles of the two views:

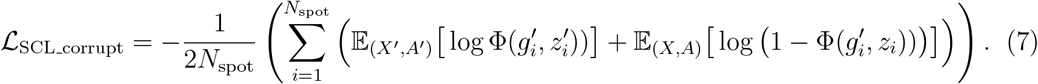

The final contrastive objective is

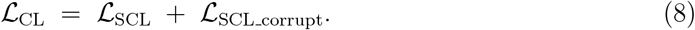

This objective maximizes agreement between each pixel and its *local* context in the original view while minimizing agreement with the corrupted view, promoting embeddings that respect spatial adjacency and discouraging spurious feature-geometry alignments.

#### 2.3.4 Training objective and optimization

The representation learning module is trained by jointly minimizing the self-reconstruction loss and the contrastive losses defined above. The overall objective is

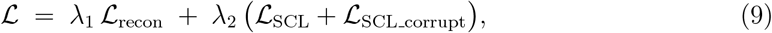

where *λ*_1_ and *λ*_2_ weight the contributions of reconstruction and contrastive terms, respectively. Unless stated otherwise, we set *λ*_1_ = 10 and *λ*_2_ = 1. We optimize Eq. (9) using the Adam optimizer[39] with a learning rate of 10^−3^ for 600 epochs.

### 2.4 Spatial domain assignment via clustering and optional refinement

After training, we obtain reconstructed spatial expression **H** ∈ ℝ^*n*×*p*^ from the decoder (Fig.1) and perform non-spatial clustering with *mclust* (Gaussian mixture modeling)[40] to assign spots to domains. Each cluster is interpreted as a spatial domain whose members share similar expression profiles and, typically, spatial proximity. For slices with manual annotations, we fix the number of clusters *K* to the annotated value. For slices without annotations, we vary *K* over a reasonable range and select the value that maximizes the average Silhouette coefficient (SC)[41] computed on **H**.

Although **H** is learned using both expression and spatial information, occasional misassignments may place isolated pixels into spatially disparate domains. To mitigate such noise, we provide an *optional* neighborhood refinement: for pixel *i*, define its local neighborhood with *r* pixels. CASTLE reassigns a pixel to the domain that is most common among its nearby spots. Empirically, *r* = 20 for spot level resolution and *r* = 50 for single-cell level resolution yielded the best performance in our benchmarks. This refinement is not recommended for ST datasets with only a few hundred spots (e.g., mouse olfactory bulb) or for high-resolution platforms such as Stereo-seq and Slide-seqV2.

### 2.5 Evaluation metric

When ground-truth labels are available, we evaluate clustering accuracy using the *Adjusted Rand Index* (ARI), which measures the agreement between predicted and true partitions while correcting for chance; ARI ranges from − 1 to 1, with larger values indicating better agreement.

When ground truth is unavailable, we report the *Silhouette Coefficient* (SC) computed on **H**. For each pixel *i*, let *a*(*i*) be the mean distance from *i* to other members of its assigned cluster and *b*(*i*) the minimum mean distance from *i* to points in any other cluster. The per-pixel silhouette is defined as:

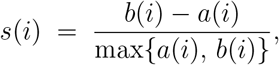

and the reported SC is the average of *s*(*i*) over all pixels; higher values indicate better separation and cohesion.

## 3 Results

In all benchmarking analyses, we estimated cell type compositions using the deconvolution pipeline (details in the Supplementary Information) and used these to construct spatial graphs for GCN-based contrastive learning.

### 3.1 CASTLE improves the identification of known layers with multicellular ST data

To evaluate spatial clustering performance, we first applied CASTLE to the 10x Visium LIBD human dorsolateral prefrontal cortex (DLPFC) dataset (55*µm* spots)[24], which comprises 12 sections with expert annotations of four or six cortical layers plus white matter (WM) derived from histology and gene markers (Fig.2b). We compared CASTLE with three state-of-the-art methods, GraphST, STAGATE, and stAA, and assessed recovery of the annotated laminar architecture based on the ARI values. Across all 12 sections, CASTLE delivered the highest median ARI together with the lowest across-section variance, indicating both better average accuracy and greater stability (Fig.2a). Although stAA achieved the highest ARI on a single section, its greater variability across sections indicates reduced robustness overall. Its median performance remained below that of CASTLE, despite generally outperforming GraphST and STAGATE. Section-level inspection supports these aggregate trends. For sample 151670 containing four layers and the white matter (WM) section, CASTLE achieved an ARI of 0.6485, whereas the other methods clustered in the 0.43–0.47 range. (Fig.2c). CASTLE recovered the major regions of all layers and WM with coherent boundaries; by contrast, GraphST tended to preserve layer ordering but split Layer 3 into two clusters, and both STAGATE and stAA over-segmented Layer 3 while failing to reliably identify WM. For sample 151673 with six layers and the WM section, CASTLE again achieved the best ARI (0.6366) and produced sharper boundaries, most notably at the Layer 1/Layer 2 interface (Fig.2d). GraphST performed secondary (ARI=0.6324) but showed less distinct separation, while STAGATE and stAA both achieved lower ARI (< 0.60). Collectively, these results indicate that CASTLE enhances laminar recovery and across-sample stability, showing clear improvements at difficult-to-separate boundaries and in WM detection compared with existing graph-based approaches.

**Figure 2:**
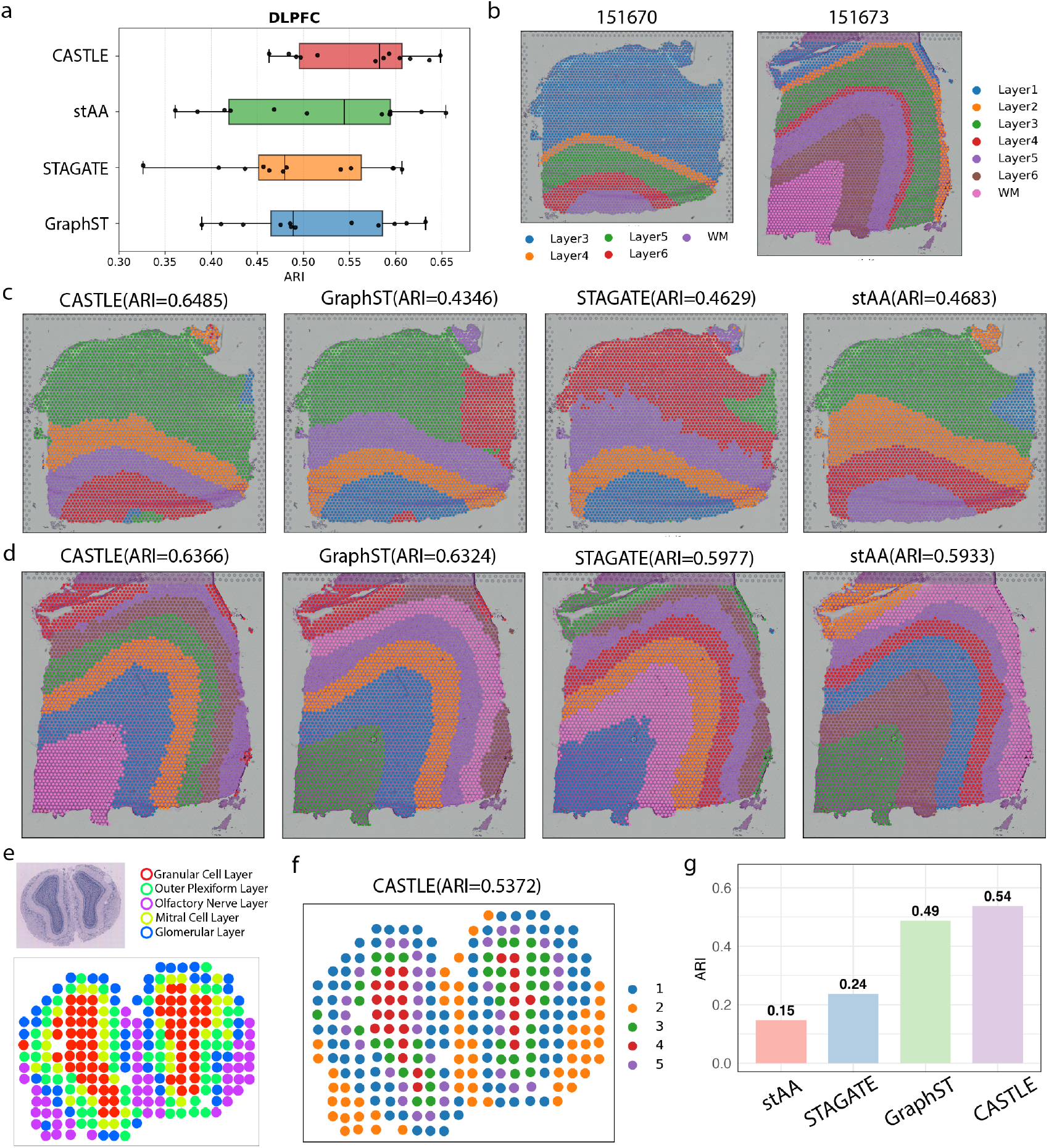
Comparison of CASTLE and other methods on ST data of different spatial resolutions. **(a)** Boxplots of adjusted rand index (ARI) scores of 12 DLPFC slices across 4 methods. **(b)** Manual annotation of the DLPFC slice 151670 and151673. **(c)** Cluster results of CASTLE, GraphST, STAGATE, and stAA for sample 151670. **(d)** Cluster results of CASTLE, GraphST, STAGATE, and stAA for sample 151673. **(e)** (Top) H&E-stained image of the mouse olfactory bulb (MOB) tissue section. (Bottom) Visualization of the corresponding MOB spots, colored by their transcriptional cluster memberships mapped to spatial locations corresponding to MOB layers from STdeconvolve[42]. **(f)** Cluster results of CASTLE in MOB. **(g)** The ARI values of four methods.

Beyond human tissue, we evaluated CASTLE on the mouse main olfactory bulb (MOB) ST data (100-µm resolution)[25]. The MOB exhibits five symmetric layers by histology, the granule cell layer (GCL), mitral cell layer (MCL), outer plexiform layer (OPL), glomerular layer (GL), and olfactory nerve layer (ONL) (Fig.2e), providing a well-defined laminar benchmark. Following prior work, we used the coarse layer annotations derived from STdeconvolve[42] as ground truth for comparison (Fig.2e). Under this setup, CASTLE accurately recovered the laminar architecture and achieved the highest ARI (0.5372) (Fig.2f). GraphST also identified layer-consistent clusters but with a lower ARI (0.4875). STAGATE and stAA produced lower ARIs (0.1468 and 0.2374, respectively) (Fig.2g); despite sometimes yielding visually sharp partitions, their clusters did not align with histological layers and frequently fragmented within layers, leading to boundaries that diverged from the known MOB organization. Overall, the results show that CASTLE more accurately captures laminar structure in lower-resolution ST data relative to composition-aware, literature-supported annotations.

### 3.2 CASTLE reveals spatial heterogeneity of human breast cancer tissue

Beyond laminar cortex, we evaluated the performance of CASTLE in resolving tumor microenvironments, which are spatially heterogeneous and mechanistically complex. We analyzed a human breast cancer 10x Visium dataset (3,798 spots, 36,601 genes) with 20 manually annotated regions (Fig.3a) grouped into four morphotypes: DCIS/LCIS (non-invasive in situ disease with an intact basement membrane), IDC (invasive ductal carcinoma breaching the basement membrane)[43], tumor edge (interface between tumor cores and adjacent stroma; typically enriched for ECM remodeling, angiogenesis and immune infiltration)[44], and healthy region. Taking the annotated 20 regions as ground truth, CASTLE achieved the best overall performance (ARI = 0.5688) (Fig.3b), with cluster patterns broadly concordant with the annotations. CASTLE merged several adjacent tumor edge regions into a single, spatially contiguous cluster, consistent with their proximity and shared biology, while further subdividing healthy tissue, IDC and DCIS/LCIS into functionally distinct subdomains that were not captured by the original coarse labels.

In the region annotated as “Healthy 1”, the enrichment guided by unique markers showed that CASTLE divided the area into two biologically distinct states rather than one section. Within this region, Cluster 4 was enriched for ECM (extracellular matrix) programs (collagen biosynthesis/organization, ECM–receptor interaction, proteoglycans) and platelet/hemostasis pathways, together with PDGF/IGF signaling, in KEGG and Reactome source pathways (Fig.3d, Fig.S4a). Expressed DCN, ACTA2, and VWF (Fig.3e, Fig.S3c), which are signatures consistent with reactive stroma, matrix remodeling, and angiogenic/platelet activity frequently implicated at tumor margins and in normal-adjacent fields [45, 46]. In contrast, Cluster 18 showed significant enrichment for oxidative phosphorylation and respiratory electron transport (Fig.3d, Fig.S4a) with CD34 expression, aligning with an endothelial/perivascular compartment that is metabolically active rather than quiescent[47, 48]. These findings indicate that histologically “healthy” tissue near tumors is not homogeneous but can encode pre-organized stromal and vascular niches (a form of field-wide change) relevant to how tumors grow, invade and respond to therapy.

**Figure 3:**
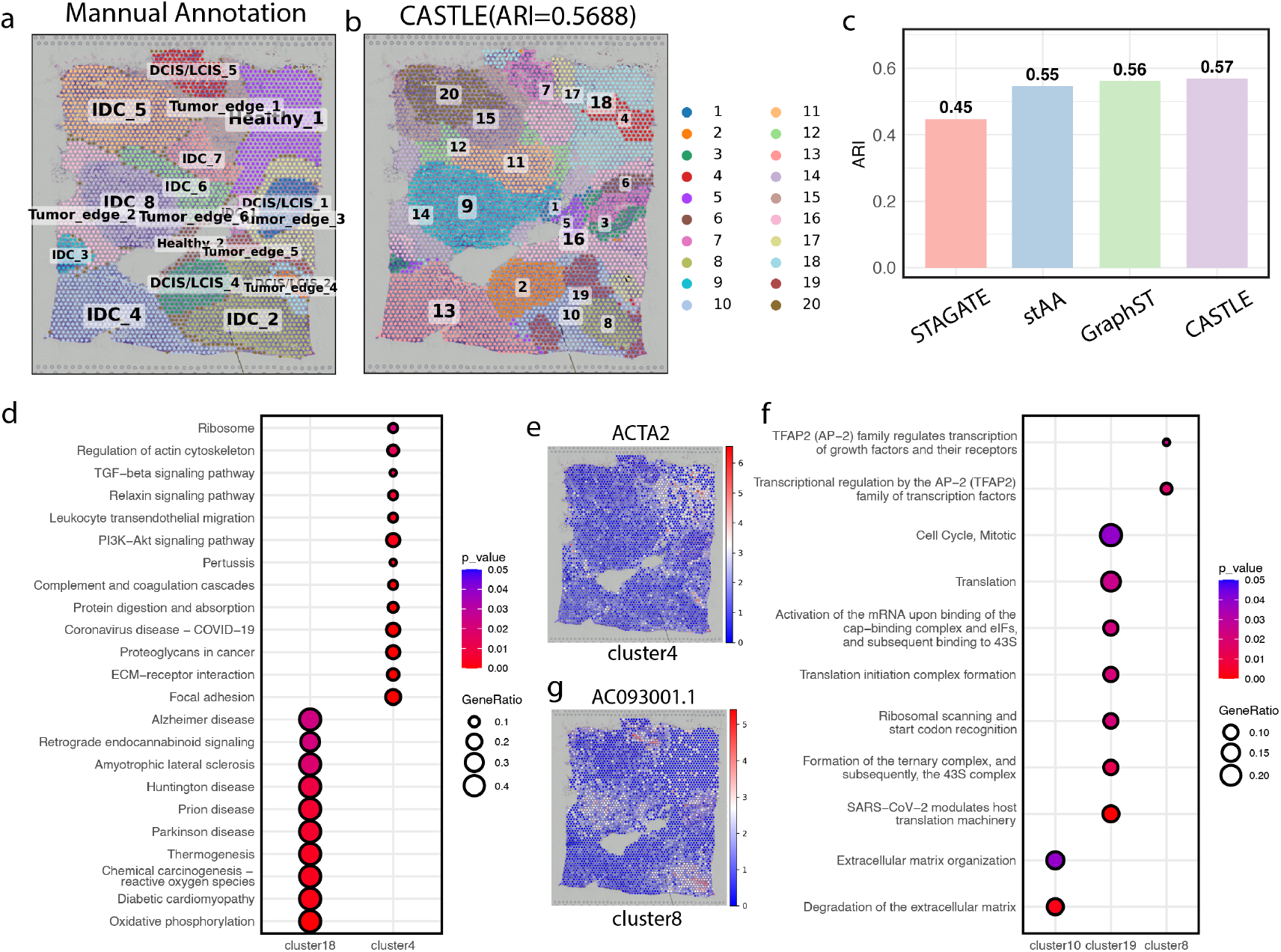
Spatial clustering on the human breast cancer ST data. **(a)** The manual domain annotation served as the ground truth. **(b)** Domains detected by CASTLE. **(c)** The ARI values of four compared methods. **(d)** The KEGG enrichment analysis of Cluster 4 and 18, focusing on the Healthy 1 region. **(e)** Expression distribution of the identified marker gene *ACTA2* associated with Cluster 4. **(f)** The comparision of Reactome enrichment analysis of Cluster 8, 10 and 19, focusing on the IDC 2 region. **(g)** Expression distribution of the identified marker gene *AC093001*.*1* associated with Cluster 8.

Within a histologically defined IDC region (“IDC 2”), CASTLE resolved three transcriptionally distinct subdomains with non-overlapping Reactome programs (Fig.3f). Cluster 8 was enriched for “Transcriptional regulation by the AP-2 (TFAP2) family,” with markers including *ESR1* and *CDH1* (Fig.S3c)), consistent with an AP-2–driven luminal/ER-positive state (TFAP2C is known to regulate ER signaling and *ESR1* expression). In this AP-2–enriched domain, the lncRNA AC093001.1 (TM4SF18-AS1) (Fig.3g) co-localized with *ESR1* and *IRF6*; while mechanistic roles remain to be tested, its spatially restricted up-regulation suggests participation in luminal regulatory circuitry[49]. Cluster 10 showed “Extracellular matrix organization” and “Degradation of the ECM,” in keeping with basement-membrane breakdown and invasion[50, 51]. Cluster 19 was dominated by cap-dependent translation initiation and mitotic cell-cycle programs, indicative of a proliferative/translation-high state[52, 53]. Thus, a single IDC region in the manual annotation actually comprises at least three mechanistically distinct tumor states pointing to different therapeutic liabilities (endocrine signaling vs. matrix remodeling vs. cell-cycle/translation control).

Finally, within DCIS/LCIS (“DCIS/LCIS 1”), CASTLE identified three subdomains that indicate early functional heterogeneity in an in-situ lesion (Fig.S4b). Cluster 3 was enriched for ribosome and multiple translation-initiation/elongation modules, consistent with a biosynthetically active epithelial state often linked to early progression[54]. Cluster 6 showed metal sequestration by antimicrobial proteins and viral mRNA translation, matching an S100A8/A9-like signature (calprotectin), which connects innate immunity, metal chelation and inflammation-associated cancer biology[55, 56]. Cluster 7 was enriched for antigen processing/presentation and ER protein-processing, together with estrogen-signaling and HSF1/HSP90 chaperone programs, its key marker *STC2* encodes a secreted glycoprotein whose expression is induced by ER stress, hypoxia and nutrient deprivation and helps tumor cells maintain redox homeostasis, avoid apoptosis and adapt to therapy[57]. Together, these DCIS/LCIS subdomains highlight mechanistically distinct epithelial and microenvironmental states—translation control, innate immune signaling, and ER-stress adaptation that are obscured by a single “in situ” label but become apparent once spatially informed clustering resolves local niches. Our results suggest that CASTLE can partition tumor tissue into functionally distinct spatial domains, providing a basis for downstream microenvironmental and pathway analyses.

### 3.3 Domian detection with cellular resolution ST data

Multiplexed error-robust fluorescence in situ hybridization (MERFISH) is a widely used imaging-based spatial transcriptomics technology that achieves single-cell resolution[58]. We assessed CASTLE on a MERFISH dataset of the mouse hypothalamic preoptic area (MPOA) comprising five annotated sections (155 genes, 15 cell types, 8 regions) (Fig.4a,b)[59]. Domain prediction is complicated here because the tissue exhibits complex organization with irregularly shaped, closely apposed regions. Leveraging the identified cell types, we constructed a cell type-aware graph and obtained strong performance (Fig.4c,d): across all five MPOA sections it achieved the highest median ARI and, aside from one outlier, outperformed the comparison methods for the remaining four sections. Because cell-type labels can be uncertain in practice, we also evaluated a label-agnostic variant (CASTLE-ES) that augments the spatial graph with gene expression similarity between cells (Fig.4d). This variant produced results competitive with CASTLE, and its median ARI exceeded those of the other compared methods, indicating improved stability (Fig.4c, Fig.S5). Together, these findings indicate that explicitly modeling local composition is advantageous when high-quality labels are available, whereas expression-similarity graphs offer a robust alternative when annotations are noisy or incomplete.

**Figure 4:**
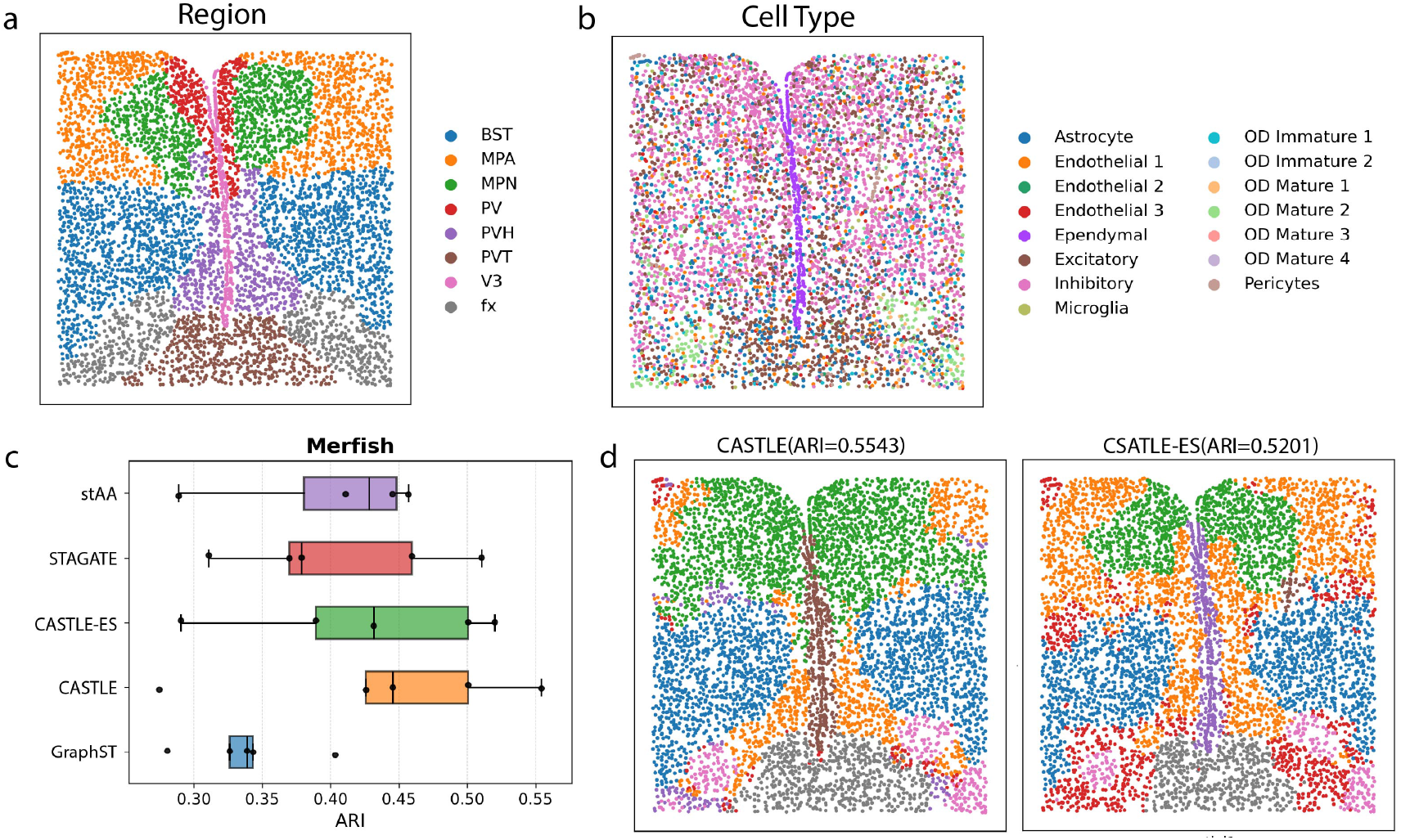
Comparison of CASTLE and other methods on MERFISH data. **(a)** The ground truth on a representative slice of MERFISH dataset. BST, bed nuclei of the strata terminalis; V3, third ventricle; PV, periventricular hypothalamic nucleus; PVT, paraventricular nucleus of the thalamus; MPN, medial preoptic nucleus; fx, columns of the fornix; PVH, paraventricular hypothalamic nucleus; MPA, medial preoptic area. **(b)** Annotated 15 cell types **(c)** Boxplots of adjusted ARI values across 5 MPOA slices with the five methods. In CASTLE-ES, we used gene expression similarity (ES) to refine the spatial graph, pretending cell type information is unavailable. **(d)** Domain detection results of CASTLE.

We further evaluated CASTLE on STARmap, an imaging-based, single-cell resolution spatial transcriptomics platform. The STARmap benchmark comprised three mouse medial prefrontal cortex sections with expert layer annotations[29]. Across methods, CASTLE and stAA achieved substantially higher ARIs than GraphST and STAGATE (Fig. S6a-b). In two sections, CASTLE’s ARI was slightly below stAA’s; in the remaining section, CASTLE clearly outperformed stAA. Overall, CASTLE delivered strong and consistent performance across all sections, whereas stAA showed higher variability.

### 3.4 CASTLE shows improved performance on subcellular ST data

Beyond commonly used single cell and multicellular resolution ST data, we further assessed the scalability and robustness of CASTLE with subcellular resolution ST technology. This inlcudes a coronal mouse olfactory bulb dataset profiled by Stereo-seq and a mouse hippocampus dataset profiled by Slide-seqV2. Stereo-seq achieves subcellular resolution using DNA-nanoball patterned arrays[34]. The DAPI-guided annotation delineates seven laminar structures, namely rostral migratory stream (RMS), granule cell layer (GCL), internal plexiform layer (IPL), mitral cell layer (MCL), external plexiform layer (EPL), glomerular layer (GL), and olfactory nerve layer (ONL) (Fig.5b). With the number of clusters fixed at seven to match the seven annotated layers, we compared CASTLE with GraphST, STAGATE, and stAA (Fig.5a). Overall, all four methods separated the outer layers, ONL, GL, and EPL. Although stAA achieved the highest SC score, only CASTLE cleanly delineated the inner structure, demarcating GCL from RMS. STAGATE failed to distinguish RMS from GCL, while GraphST and stAA frequently mixed GCL with IPL. Consistent with these layer patterns, domain marker evaluation showed good correspondence between CASTLE’s domain marker genes and the expected layer identities (Fig.5c).

Slide-seqV2 profiles expression at higher spatial density (10,000 beads per section), though fixed bead locations may capture mixtures of nearby cells[35]. We applied CASTLE to the Slide-seqV2 mouse hippocampus dataset[27] with a cell type composition-aware graph that accounts for multiple cell types per bead. Using the Allen Brain Atlas annotations as ground truth, we evaluated recovery of hippocampal regions (CA1/CA2, CA3, DG) (Fig.S7b)[60]. STAGATE did not distinguish CA1 from CA3, obscuring the hippocampal structure. By contrast, CASTLE, GraphST, and stAA all identified CA1, CA3, and DG, though GraphST showed a cluster that mixed hippocampus with ventricular systems/epithalamus. Using the silhouette-based SC metric, stAA yielded the highest SC, likely reflecting its reliance on pre-clustering to refine embeddings, while CASTLE exceeded GraphST and STAGATE (Fig.S7a). At the gene level, CASTLE recovered canonical markers, Wfs1 (CA1), Ly6e (CA3), and C1ql2 (DG), supporting the anatomical assignments (Fig.S7c).

**Figure 5:**
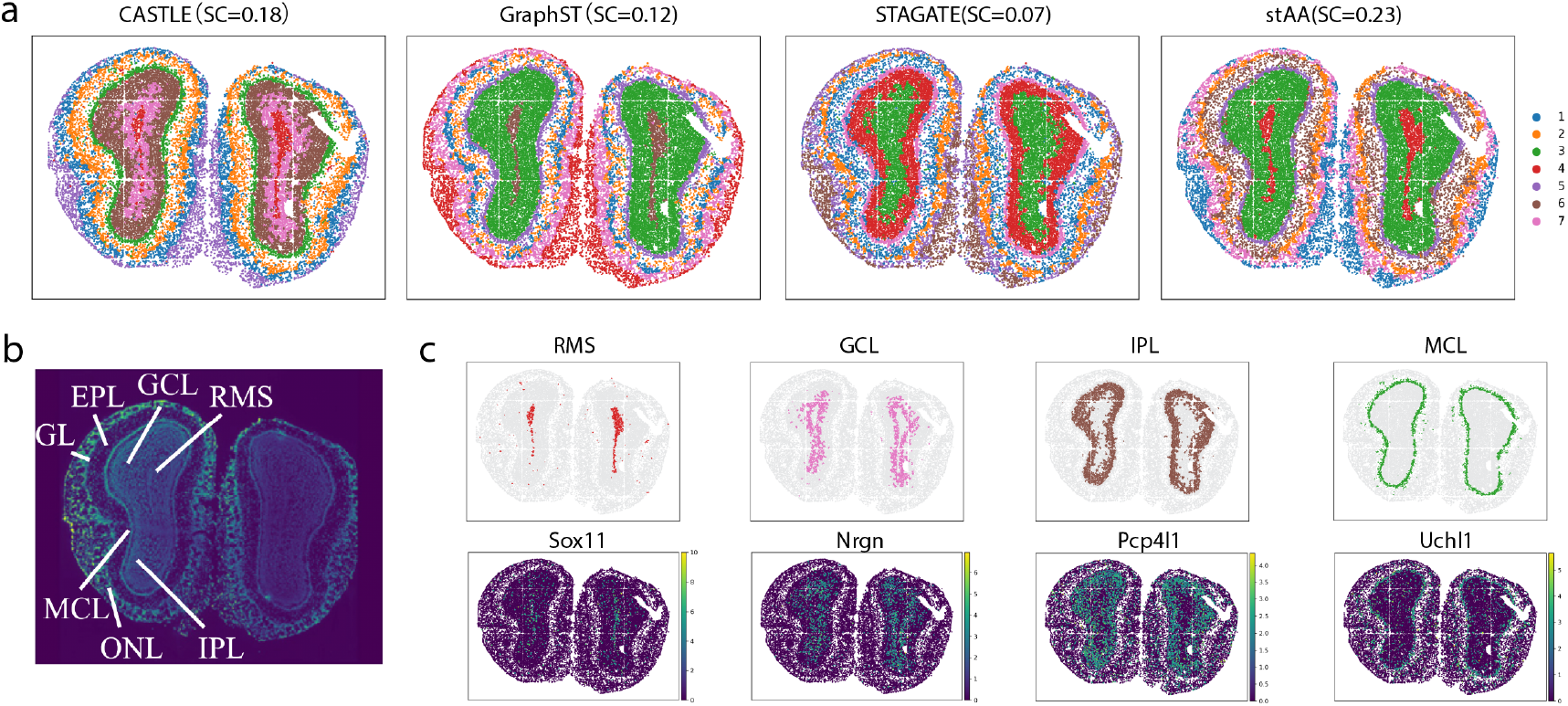
Comparison of CASTLE and other methods on Stereo-seq data. **(a)** Spatial domain identified by CASTLE, GraphST, STAGATE, and stAA in the MOB Stereo-seq data. **(b)** Laminar organization of MOB annotated in the DAPI-stained image. **(c)** Visualization of the spatial domains identified by CASTLE and the corresponding identified marker gene expressions.

## 4 Discussion

Accurate identification of spatial domains and subsequent discovery of domain marker genes is essential for understanding tissue organization and function. We presented CASTLE, a cell type-aware graph learning framework that integrates spatial proximity and spatial cell type composition information via contrastive learning embedding for spatial domain detection. A lightweight GCN encoder–decoder reconstructs expression, and a local-context contrastive objective aligns embeddings to their immediate neighborhoods and reduces spurious feature-spatial correlations. In doing so, CASTLE addresses a central challenge in spatial clustering: constructing graphs that capture both physical adjacency and biologically meaningful similarity, while providing a principled fallback to expression-based similarity when spatial labels are noisy or missing.

CASTLE introduces three key advances. First, it constructs a biologically grounded, cell-type-aware spatial graph that yields neighborhoods reflecting true tissue organization without relying on circular pre-clustering. Second, its local-context contrastive objective aligns each spot with its immediate microenvironment while reducing spurious feature-spatial correlations, producing sharper and more reliable domain boundaries. Third, CASTLE provides a unified, lightweight framework that adapts naturally across spot-, cell-, and subcellular-resolution platforms. Together, these innovations enable CASTLE to generate robust, interpretable, and biologically meaningful spatial domains across diverse technologies.

We evaluated CASTLE against state-of-the-art methods[6] on various ST platforms and resolutions. Consistent gains appeared in both accuracy and stability. On 10x Visium DLPFC, CASTLE achieved the highest median ARI and the lowest variance, recovering challenging layer boundaries and improving the white matter clustering. In mouse MOB ST data, it separated inner layers GCL vs. RMS, that were conflated by competing methods. In a breast cancer 10x Visium dataset, it attained the best overall ARI and uncovered functionally distinct subdomains within healthy, IDC, and DCIS/LCIS regions; enrichment analyses of cluster-unique markers highlighted distinct pathway programs in each compartment. At single-cell resolution MERFISH MPOA data, the cell type–aware graph variant yielded the best median ARI, while the label-free expression-similarity graph remained competitive and more stable than the counterparts when labels were uncertain. On STARmap, CASTLE matched or exceeded state-of-the-art performance with lower variability. Analyses on Stereo-seq and Slide-seqV2 further demonstrated scalability and anatomical fidelity, including recovery of multiple layers and canonical markers.

Incorporating cell type information into spatial graph construction is crucial for capturing biologically meaningful tissue organization and improving domain detection accuracy. Two methodological choices are central to these outcomes. First, cell type-aware edges refine the spatial graph by suppressing cross-type connections that arise from spatial proximity alone, thereby sharpening domain boundaries, especially when spots contain mixed cell populations or when neighboring regions differ mainly in composition. Second, local contrastive learning with a corrupted view encourages embeddings to align with their true local context while penalizing spurious feature-geometry correlations, leading to improved generalization across spatial transcriptomics platforms.

Although CASTLE performs robustly across diverse platforms, several limitations remain. The method assumes that spatial domains are at least partly reflected in local cell-type composition; in tissues where boundaries are driven primarily by transcriptional state or microenvironmental cues, composition-aware graphs may provide limited benefit. In settings dominated by continuous gradients rather than discrete compartments, CASTLE may smooth transitions or merge adjacent regions with similar mixtures. Because CASTLE emphasizes composition-informed neighborhoods, it may also merge domains that share cell types but differ in finer functional states unless these differences are strongly encoded in expression. In addition, inaccuracies in deconvolution, cell-type annotation, or spatial registration can propagate into graph construction and affect clustering. Future extensions that incorporate uncertainty estimates, histology-derived features, or multi-slice integration may help address these challenges.

In summary, CASTLE is a practical and versatile approach for spatial domain identification that attains finer-grained, biologically interpretable structure across spot-, bead-, and single-cell-resolution datasets. Remaining limitations include dependence on deconvolution or existing annotations when available, which may propagate labeling errors, sensitivity to graph hyperparameters at extreme resolutions, and a current focus on single-sample graphs without explicit batch integration. Future work could incorporate histology-derived features and uncertainty estimates into edge weighting, and extend the framework to multi-slice or multi-batch graphs through domain adaptation and joint deconvolution-clustering.

## Author contribution

Y.C. conceived the idea. A.X. and Y.C. designed the experiments. A.X. developed the method, implemented the software, performed data analysis. A.X. and Y.C. wrote the manuscript.

## Conflict of interest

None declared.

## Data availability

All data analyzed in this study are available in their raw form from the original sources. Specifically, the DLPFC dataset[24] is accessible within the spatialLIBD package (http://spatial.libd.org/spatialLIBD). Data and H&E images for MOB[25] are available for download at https://www.spatialresearch.org/resources-published-datasets/doi-10-1126science-aaf2403/. The human breast cancer 10x Visium dataset[33] is accessible on https://www.10xgenomics.com/resources/datasets/human-breast-cancer-block-a-section-1-1-standard-1-1-0 The MERFISH dataset[58] of the mouse hypothalamic preoptic area (MPOA) and mouse medial prefrontal cortex data from STARmap[29] are available at http://sdmbench.drai.cn/. The Slide-seq dataset[27] is available at https://portals.broadinstitute.org/single_cell/study/slide-seq-study. The processed Stereo-seq data from mouse olfactory bulb tissue[34] is accessible on https://github.com/JinmiaoChenLab/SEDR_analyses.

## Code availability

Python codes to implement CASTLE are available in the GitHub repository at https://github.com/Cui-STT-Lab/CASTLE

## Notes

### Competing Interest Statement

The authors have declared no competing interest.

