## Supplementary Materials for "CASTLE: Cell-type Aware SpaTial domain detection via contrastive Learning Embedding"

### Supplementary material for "CASTLE: Cell-type Aware SpaTial domain detection via contrastive Learning Embedding"

#### 1 Description of real datasets and cell type deconvolution

For 10x ST data, which contains multiple cells per spot, we implemented RCTD to obtain the cell type composition for each pixel. We then set $k = 5$ and the threshold $\tau = 0.9$ to control the average number of neighbors when constructing the spatial graph for fitting the GCN model.

##### 1.1 DLPFC 10x Visium data

The DLPFC dataset contains an average of 3,499 spots and 33,538 genes. This dataset is released on the 10x Genomics Visium platform and well well-annotated by the spatialLIBD project. Five or seven domains are manually labeled on these samples, and each region has a clear boundary. We applied RCTD [1] to all 12 samples using a single-cell reference [2] comprising seven major cell types.

##### 1.2 MOB ST data

We obtained mouse olfactory bulb (MOB) datasets from the original publication, focusing on MOB replicate #8, with 260 spots and 14,828 genes. Coarse clustering annotations were obtained from STdeconvolve. The reference scRNA-seq data is obtained from the Gene Expression Omnibus (GSE121891[3]) with eleven major cell types.

##### 1.3 Human breast cancer

The human breast cancer ST data contains 3,798 spots and 36,600 genes. The authors in SEDR[4] manually labeled this data, relying on the morphological image and gene expression profiling. Twenty domains are recognized in this dataset, including four morphotypes which are ductal carcinoma in situ/lobular carcinoma in situ (DCIS/LCIS), invasive ductal carcinoma (IDC), tumor edge areas, and healthy regions. The scRNA-seq datasets of breast cancers used here are publicly available from the Gene Expression Omnibus database GSE176078[5] with thirteen major cell types.

##### 1.4 Merfish MPOA data

The MERFISH dataset is an imaging-based ST dataset published in 2018[6]. Among all slices, five slices were annotated with region labels from BASS[7]. The numbers of cells are 5,488 (Slice

0.04), 5,557 (Slice 0.09), 5,926 (Slice 0.14), 5,803 (Slice 0.19) and 5,543 (Slice 0.24). The number of unique genes is 155. This dataset with annotation was downloaded from SDMBENCH[8].

In the CASTLE-ES, we obtained cosine similarity of gene expressions and set the threshold $\tau = 0.6$ in the nearest 10 neighbors to refine the edge of the spatial graph.

##### 1.5 STARmap data

STARmap utilizes combinatorial barcoding and imaging to detect transcriptome-wide expression patterns at single-cell resolution in situ[9]. This dataset contains three slices from the mouse medial prefrontal cortex, with expert annotations of layers[7]. The numbers of cells are 1,049 (Slice 0.69), 1,053 (Slice 0.77) and 1,088 (Slice 0.70). The number of unique genes is 166. This dataset with annotation was downloaded from SDMBENCH[8].

Here, we utilized the cosine similarity in the neighbor to construct the graph, rather than using the cell type information. The mouse medial prefrontal cortex is roughly partitioned into four laminae, whereas the STARmap dataset profiles $\sim 1,000$ cells annotated to fifteen finer cell types. Injecting these fine-grained labels directly into the graph can introduce confounding. For example, Layer 1 is enriched for both smooth muscle cells (SMC) and endothelial cells (Endo); although both reside within the same layer, their spatial distributions differ. Hard, label-driven edges may therefore connect SMC–SMC or Endo–Endo across microdomains and artificially split a single anatomical layer, leading to over-segmentation and misclustering. A softer, composition-aware, or expression-similarity approach is preferable in this setting.

##### 1.6 Stereo-seq MOB data

The Stereo-seq dataset is obtained by high-resolution full-transcriptome coverage technologies (Stereo-Seq technology). It contains 19,109 pixels and 27,106 genes. Given its spatial resolution of $14\mu$m, we first applied RCTD to perform cell-type deconvolution using the same single-cell reference as in the ST MOB dataset. We then set $k = 4$ and the threshold $\tau = 0.4$ to control the average number of neighbors in the GCN model.

##### 1.7 Slide-seqV2 hippocampus data

Slide-seqV2 mouse hippocampus and related single-cell reference were downloaded from RCTD[1] directly. We applied RCTD to do the deconvolution with 17 cell types, and then set $k = 6$, with threshold $\tau = 0.5$ to control the average number of neighbors for the model.

### 2 Supplementary Figures

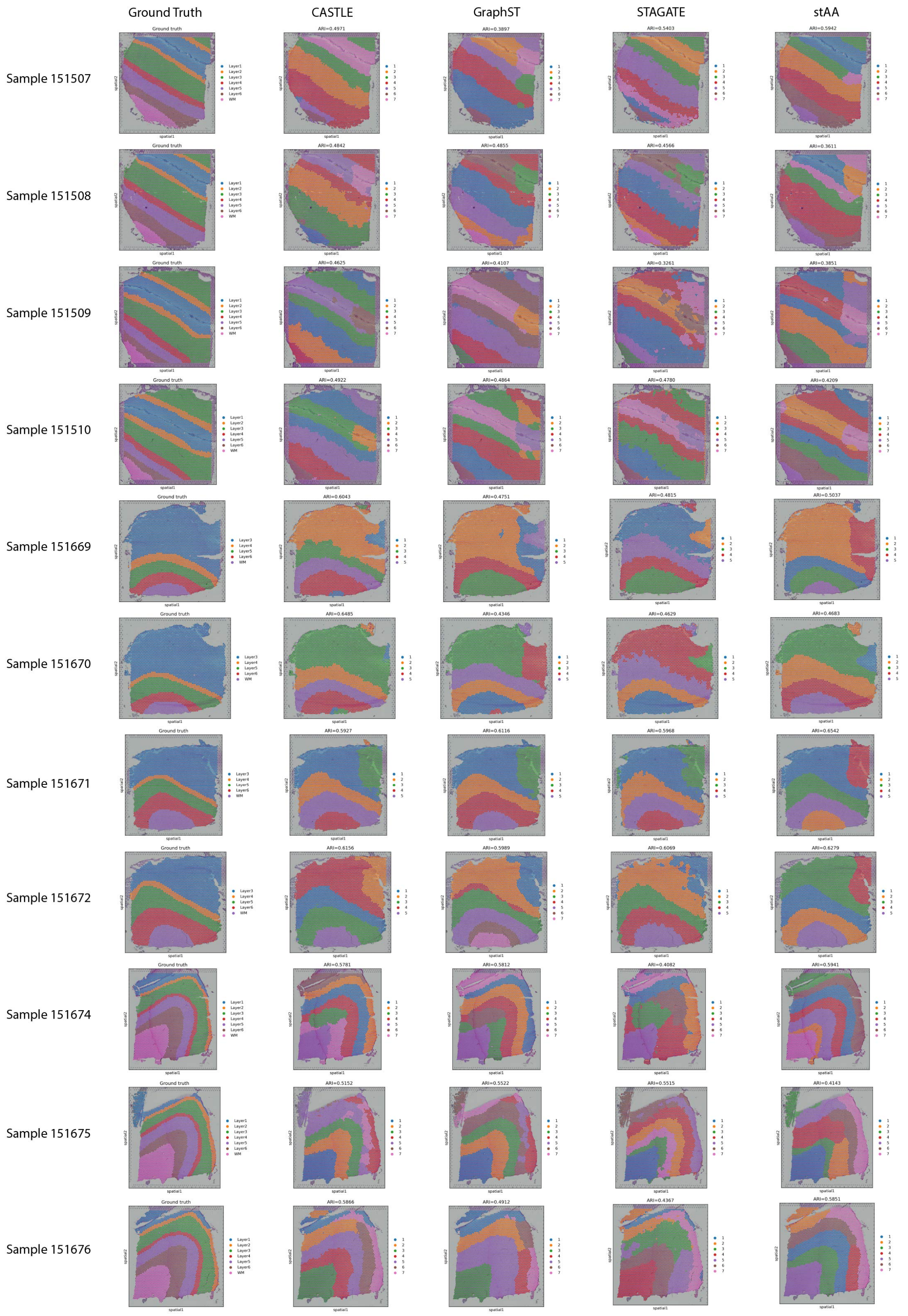


Figure S1: **Domain detection results from CASTLE and other methods on 11 DLPFC slices.**

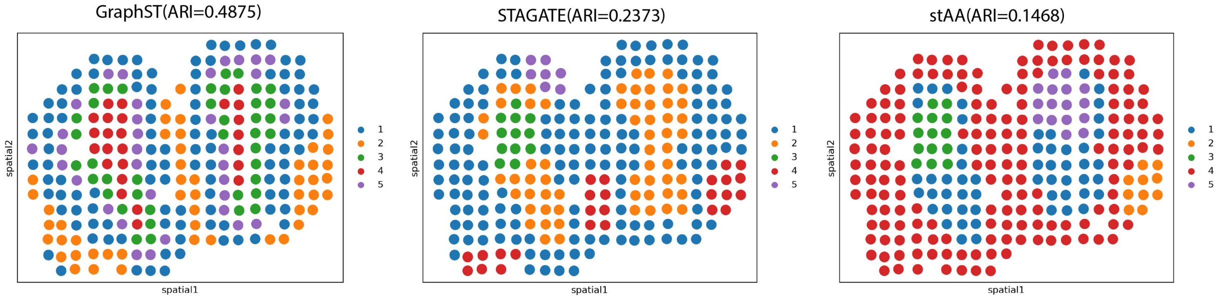


Figure S2: **GraphST, STAGATE and stAA clustering results on MOB ST data.**

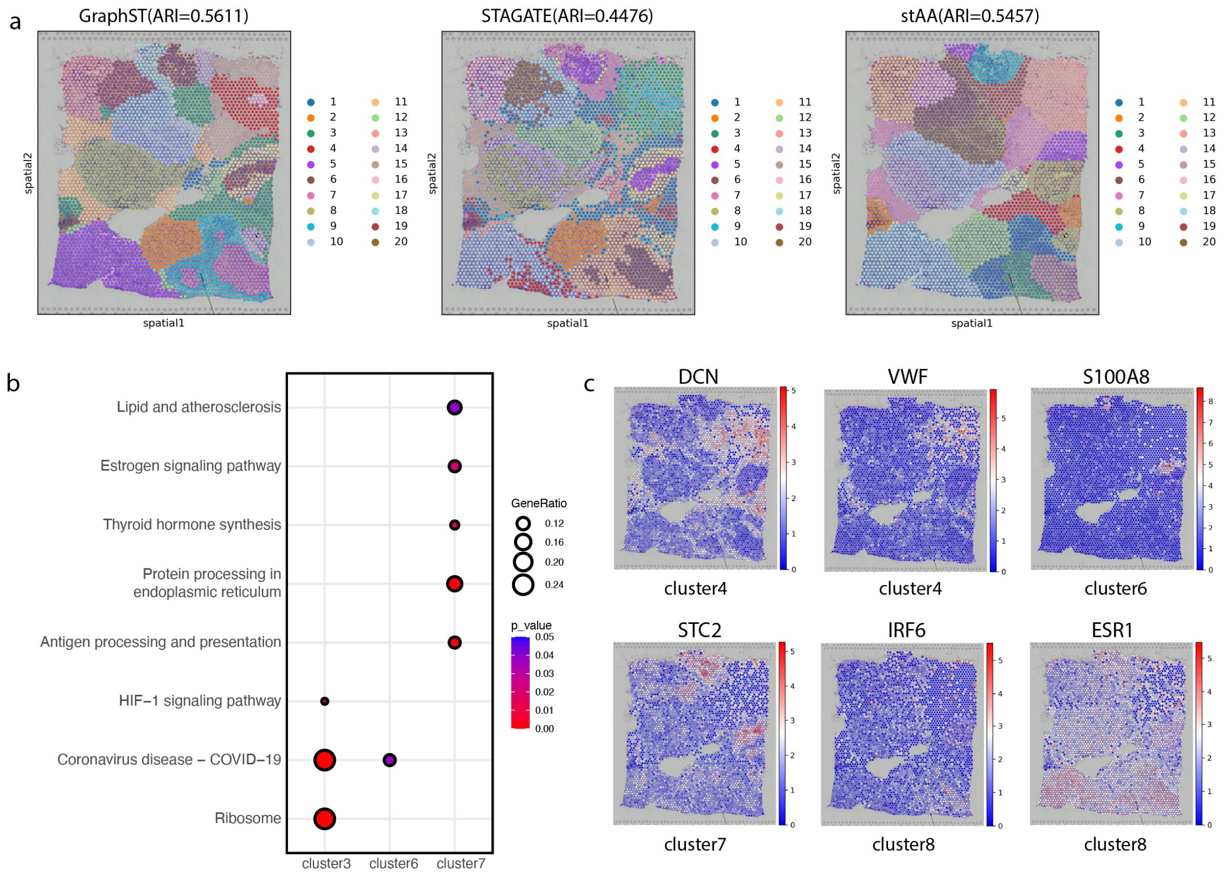


Figure S3: **Clustering results of the human breast cancer 10x data.** **(a)** Clustering results of GraphST, STAGATE and stAA. **(b)** KEGG enrichment analysis of cluster3, cluster6 and cluster7, which were identified in the DCIS/LCIS region. **(c)** Expression distribution of domain marker genes.

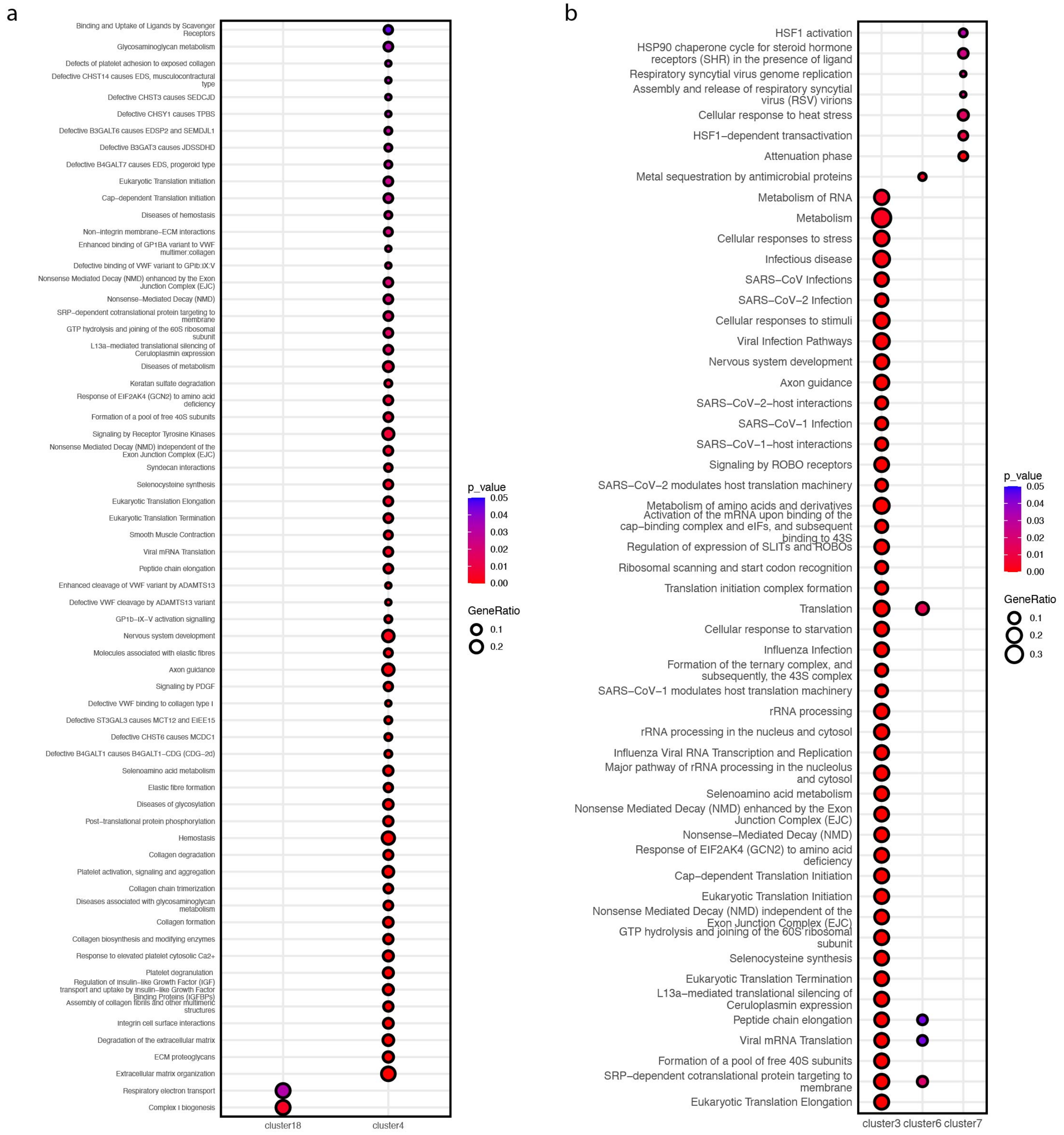


Figure S4: **Reactome enrichment analysis in human breast cancer data (a)** Reactome enrichment analysis of cluster4 and cluster18, which were identified in the healthy region. **(b)** Reactome enrichment analysis of cluster3, cluster6 and cluster7, which were identified in the DCIS/LCIS region.

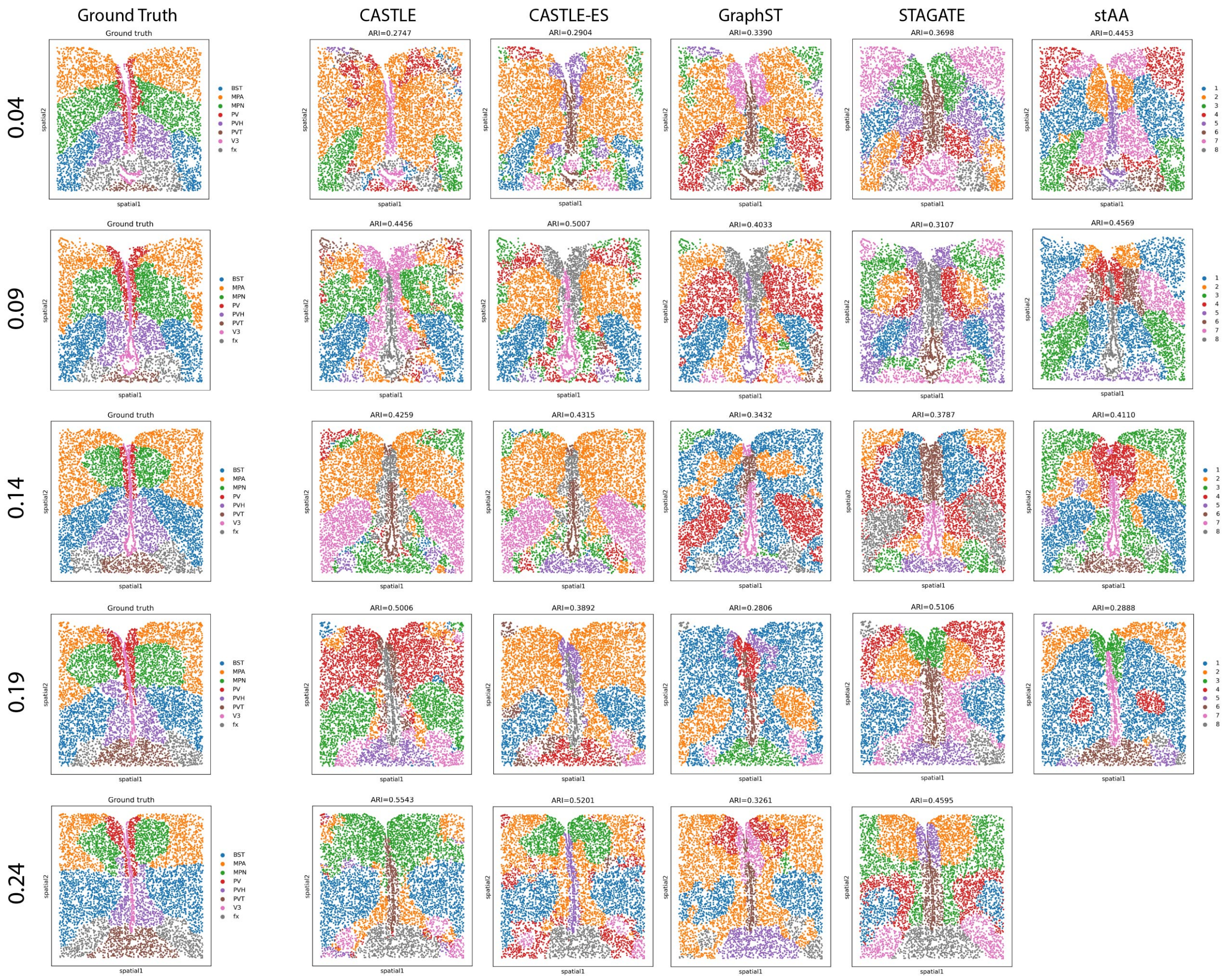


Figure S5: **Comparison between CASTLE and other methods on MERFISH MPOA data.** Clustering results across 5 tissue slices. stAA can not work on slice 0.24. The five slice samples are labelled as 0.04, 0.09, 0.14, 0.19 and 0.24.

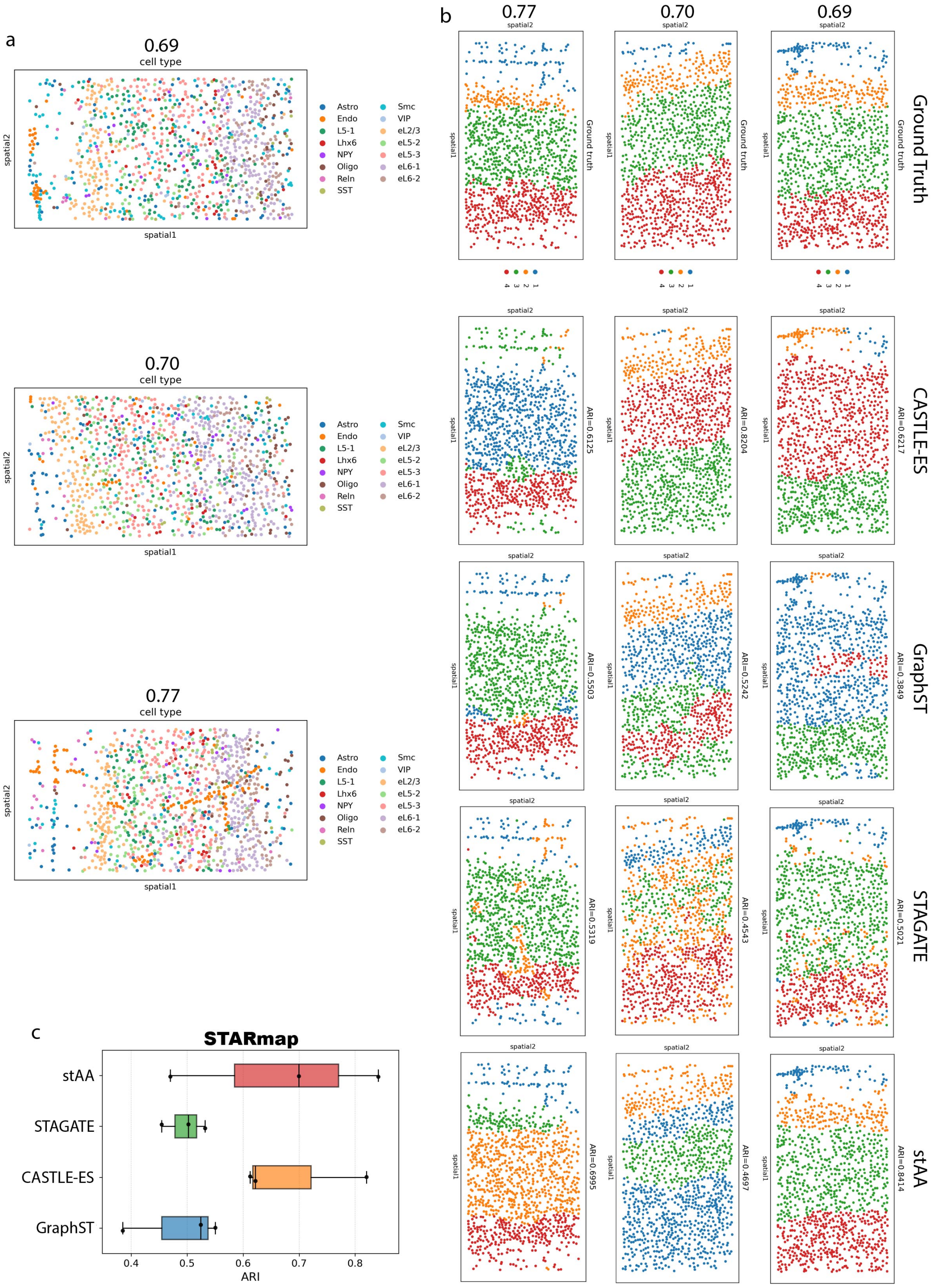


Figure S6: **Comparison between CASTLE and other methods on STARmap mouse medial prefrontal cortex data.** **(a)** Cell types spatial patterns in STARmap mouse medial prefrontal cortex data. **(b)** Clustering results of CASTLE-ES, GraphST, STAGATE and stAA. **(c)** Boxplots of adjusted ARI values across the four methods. The three samples are labelled as 0.69, 0.70 and 0.77.

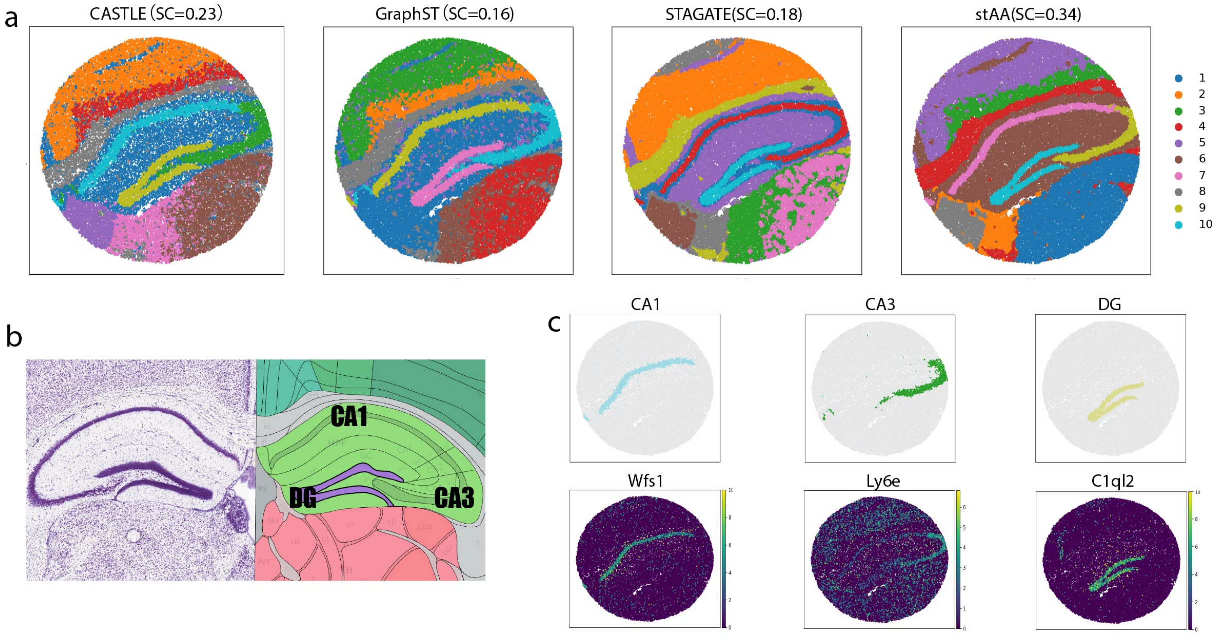


Figure S7: **Comparison between CASTLE and other methods in the mouse hippocampus Slide-seqV2 data.** **(a)** Clustering results of CASTLE, GraphST, STAGATE and stAA. **(b)** Histological annotation regions[10]. **(c)** CASTLE identified matching regions and associated domain marker genes.
